# Adaptation to sorbic acid in low sugar promotes resistance of spoilage yeasts to the preservative

**DOI:** 10.1101/2023.05.30.542886

**Authors:** Harry J. Harvey, Alex C. Hendry, Marcella Chirico, David B. Archer, Simon V. Avery

## Abstract

The weak acid sorbic acid is a common preservative used in soft drink beverages to control microbial spoilage. Consumers and industry are increasingly transitioning to low-sugar food formulations, but potential impacts of reduced-sugar on preservative efficacy are barely characterised. In this study, we report enhanced sorbic acid resistance of spoilage yeasts in low-glucose conditions. We had anticipated that low glucose may induce respiratory metabolism, previously shown to be targeted by sorbic acid. However, a shift from respiratory to fermentative metabolism was correlated with the sorbic acid resistance in low glucose. Fermentation-deficient yeast species did not show the low-glucose resistance phenotype. Phenotypes observed for certain yeast deletion strains suggested roles for glucose signalling and repression pathways in the sorbic acid resistance at low glucose. This low- glucose induced sorbic acid resistance was alleviated by supplementing yeast cultures with succinic acid, a metabolic intermediate of respiratory metabolism (and a food-safe additive) that promoted respiration. The results indicate that metabolic adaptation of spoilage yeasts promotes sorbic acid resistance at low glucose, providing new insight into potential spoilage, and preservation, of foodstuffs as both food producers and consumers move towards a reduced-sugar landscape.

## Importance

In recent years, driven by factors such as government legislation and public health concerns, consumers and the industry have been migrating to lower sugar food or drink formulations. However, the potential impacts of low-sugar formulations on preservative sensitivity of spoilage organisms has barely been addressed. In this work, we show that there is a marked impact of glucose concentration on the response of spoilage yeasts to the preservative sorbic acid, giving elevated resistance at low glucose. Furthermore, we demonstrate a novel mechanism of preservative resistance (a shift from respiratory to fermentative metabolism) and a potential method to mitigate this resistance, by addition of food safe succinic acid. This work highlights how food preservation practices may need to develop in tandem with the migration to reduced sugar foods, and also gives significant new insights to spoilage-yeast physiology.

## Introduction

Annual soft drinks consumption by adults in Europe is reportedly ∼40 litres per capita (Walton and Wittekind, 2023). Traditionally, soft drink beverages tend to contain relatively high sugar concentrations (up to 14% w/v), have a pH range of 2.4–5.0, and are commonly stored at ambient temperatures (Shankar et al., 2021). These conditions are conducive for fungal growth and, therefore, soft drinks are prone to spoilage by filamentous fungi or yeasts, with *Zygosaccharomyces bailii* being among the most prominent of spoilage yeasts. To mitigate spoilage, the weak acid (WA) sorbic acid is a common preservative used in soft drink beverages, having been widely shown to possess strong antifungal properties (Plumridge et al., 2004, Sofos and Busta, 1981, Stratford et al., 2013). In more recent years, consumer concerns over high-sugar foods and implementation of sugar tax levies in countries such as the UK, has led many beverage companies to transition towards low-sugar drinks formulations (WHO, 2022). However, potential impacts of such reformulation on preservative resistance and spoilage propensity have yet to be fully explored. Considering that nutrient (e.g. sugar) availability is well known to affect microbial physiology, e.g., nutrient assimilation rates (Schreiber et al., 2016), mitochondrial morphology (Bagamery et al., 2020), and general stress response (Causton et al., 2001, Kuang et al., 2017), there is strong rationale for anticipating impacts of altering food formulation in this way on the propensity for spoilage.

The concern raised above may be especially pertinent to spoilage by yeasts, as many yeasts are known to switch from fermentative to respiratory metabolism as extracellular glucose levels are decreased (Kayikci and Nielsen, 2015) and sorbic acid has been shown to exert greater inhibition of cells growing on respiratory substrates (e.g. glycerol) than fermentative substrates (high glucose) (Stratford et al., 2020). The latter was related to observations in the same study that the weak acid selectively targets respiration. This evidence suggested that a bias to respiratory metabolism by spoilage yeasts in low-sugar beverage conditions would sensitise them to inhibition by sorbic acid.

In this study, we tested this hypothesis by investigating effects of sorbic acid on several spoilage-yeast species in low glucose conditions, giving new insights to the relationship between carbon metabolism and preservative resistance.

## Methods

### Yeasts and Culture Conditions

The yeast species and strains used in this study were: *Saccharomyces cerevisiae* W303 (MAT**α** ura3-1 ade2-1 trp1-1 his3-11,15 leu2-3,112); *S. cerevisiae* BY4743 (MAT**a**/MAT**α** his3Δ 1/his3Δ 1 leu2Δ 0/leu2Δ 0 met15Δ 0/MET15 LYS2/lys2Δ 0 ura3Δ 0/ura3Δ 0) and its isogenic deletion strains (purchased from EUROSCARF, Frankfurt); *Zygosaccharomyces bailii* NCYC 1766 (Stratford et al., 2020); *Brettanomyces bruxellensis* (an isolate from a food factory in the UK); and fermentation deficient strains *Rhodotorula mucillaginosa* CMCC2663, *Rhodotorula glutinis* NCYC59, *Rhodotorula glutinis* (an isolate from a food factory in Israel), *Cryptococcus laurentii* (an isolate from a food factory in Rio de Janeiro, Brazil) and *Cryptococcus magnus* (an isolate from a food factory in Samara, Russia). Yeasts were maintained and grown in YPD medium [2% w/v peptone (Oxoid), 1% w/v yeast extract (Oxoid), 2% w/v D-glucose]. Where required, medium was solidified with 2% (w/v) agar. For starter cultures, single colonies were used to inoculate 10 ml of medium in 50 ml Erlenmeyer flasks and incubated with orbital shaking (New Brunswick Scientific) at 120 rev min^−1^ at 24 °C. Overnight cultures were then used to inoculate fresh medium to an OD_600_ 0.5 before incubating for ∼4 h as described above to allow cells to reach exponential phase.

### Growth Assays

Growth assays were conducted in 96-well microtiter plates (Greiner), with each well containing 100 µl YP [2% peptone (Oxoid), 1% yeast extract (Oxoid)] pH4.0 (adjusted with HCl)] supplemented with either 2% or 0.1% (v/w) D-glucose. For addition of weak acids, aliquots from stock solutions were added to each well to achieve the desired final acid concentration. Stock solutions of sorbic acid, propionic acid and decanoic acid were dissolved in water. Acetic acid was dissolved in YP to prevent the substantial dilution of culture medium by the large volumes of acetic acid needed. Typically, final weak acid concentrations were ∼55–60% of the MIC. Wells were inoculated with exponential-phase cells to a final optical density (OD_600_) of 0.1. Growth was monitored during incubation at 24 °C in a BioTek Powerwave XS microplate spectrophotometer, shaking for 2 min before OD_600_ measurements every 30 min.

For experiments in anaerobic conditions, growth assays were conducted as above, with the exception that plates were incubated in an anaerobic chamber [Whitley DG250 anaerobic workstation; Don Whitley Scientific (10% CO2, 10% H2, 80% N2)] for 5 h to remove oxygen from culture media before inoculation of cells, as above, within the chamber to a final OD_600_ 0.1. Growth yield was assessed by OD_600_ readings after 30 h incubation in the anaerobic chamber at 24 °C.

### Intracellular pH Measurements

Prior to intracellular pH measurements, exponential phase cells were incubated for 4 h with shaking at 120 rev min^−1^, 24 °C in 10 ml of medium in 50 ml Erlenmeyer flasks, containing the weak acids and carbon sources as described in Sup. Fig. 2. Intracellular pH was measured with a method adapted from (Stratford et al., 2014). Cells were harvested from 1 ml of culture at OD_600_ ∼0.5 in each experimental condition by centrifugation at 3000 *g*, 4 min and resuspended in 1 ml phosphate buffered saline (PBS) (137 mM sodium chloride, 2.7 mM potassium chloride, 11.9 mM phosphate buffer) containing the pH sensitive intracellular stain 5-(and-6)-carboxyfluorescein diacetate, succinimidyl ester (CFDA-SE) at a final concentration of 5 mg/L. Cells were incubated in the CFDA-SE solution for 20 minutes at 24 °C, before centrifugation at 4000 *g*, 3 min and aspiration of the staining solution. Stained cells were then resuspended in the spent medium that they had been harvested from (medium was filter sterilised beforehand using a 0.22 µm filter) and incubated for 1 hr before examination by flow cytometry, described below.

A standard curve for intracellular pH was produced by staining exponential phase cells as above, followed by 1 h incubation in permeabilization solution (PBS with 100 µm nigericin) that had been pH-adjusted between 4.0–7.5 in pH 0.5 increments, using HCL or NaOH. This procedure yields cells with defined intracellular pH (Lin et al., 2003). These were analysed by flow cytometry as described below. The resultant standard curve was used to convert ratiometric fluorescence values from experimental cultures to median intracellular-pH values.

CFDA-SE ratiometric fluorescence was determined for 10^6^ cells per sample by flow cytometry, with a FACSCanto A (BD Biosciences) instrument. Laser excitation was at 488 nm and emission was collected through a 530/30 nm filter for pH-dependent emission and a 585/42nm filter for pH-independent emission from CFDA-SE. Events (cells) were gated by forward scatter and side scatter to exclude doublets and debris. Median fluorescence of gated cells was then calculated using Kaluza Analysis V2.1 software.

### Fermentation Measurements

To measure pressure resulting from gas evolution during fermentation, McCartney bottles containing 5 ml YP (supplemented with glucose as specified in the Results) were inoculated with exponential phase cells to a final OD_600_ 0.2. Bottles were then sealed with hole-punched metal caps containing intact rubber seals to prevent gas leakage. After static incubation for either 1 or 7 days at 24 °C, pressure accumulated from culture gas evolution was measured using a 2000P Differential Manometer (Digitron), alongside culture OD_600_ measurements, as above. Pressure readings were subsequently normalised to OD_600_ readings.

### Oxygen Depletion Measurements

To measure oxygen depletion from culture media as a result of respiration, McCartney bottles were each fitted with a single Planar Oxygen-Sensitive Spot (PreSens, Regensburg). These spots are coated with an oxygen-quenching fluorescent material, which can both be non-invasively excited, and emission (oxygen-dependent) read, using a FitBox 4 (PreSens, Regensburg). YP medium (5 ml), supplemented with glucose as described in Results, was added to the McCartney bottles and inoculated with exponential phase cells to a final OD_600_ 0.5. Bottles were then sealed before baseline oxygen measurements were taken, followed by static incubation for 2 h at 24 °C. After incubation, endpoint readings were taken to determine oxygen depletion from the medium as a result of respiration. Although oxygen depletion occurred over a 2 h period (Sup. Fig. 6), 1 h incubation was sufficient to measure oxygen depletion from the medium before substantial culture growth occurred and 1 h was used as the endpoint for subsequent assays.

## Results

As recent work has indicated that the food preservative sorbic acid targets respiration in yeasts (Stratford et al., 2020), we tested the hypothesis that the greater relative respiratory (versus fermentative) metabolism of yeasts at low glucose may sensitize them to sorbic acid. Yeasts were cultured with either 2% w/v glucose (∼111 mM, a standard concentration in media) or 0.1% w/v glucose (∼5.55 mM, the minimum concentration found to permit reproducible batch growth) in the absence or presence of sorbic acid. Unexpectedly, sorbic acid was less inhibitory to growth of key spoilage yeasts (*Zygosaccharomyces bailii, Saccharomyces cerevisiae, Brettanomyces bruxellensis*) at 0.1% glucose than at 2% glucose, relative to the respective corresponding controls without sorbic acid (Figure 1A, C, E). This effect could be visualised more clearly after normalising the OD values with sorbic acid to the corresponding control ODs, for each glucose condition (Figure 1B, D, F). Nevertheless, in the case of *Saccharomyces cerevisiae* W303 between ∼20 to ∼50 hours, there was outright greater growth with sorbic acid at low- versus high-glucose regardless of controls (Figure 1C) (after which time cells in high glucose began to outgrow those in low glucose, possibly due to excess remaining glucose).

**Figure 1.**
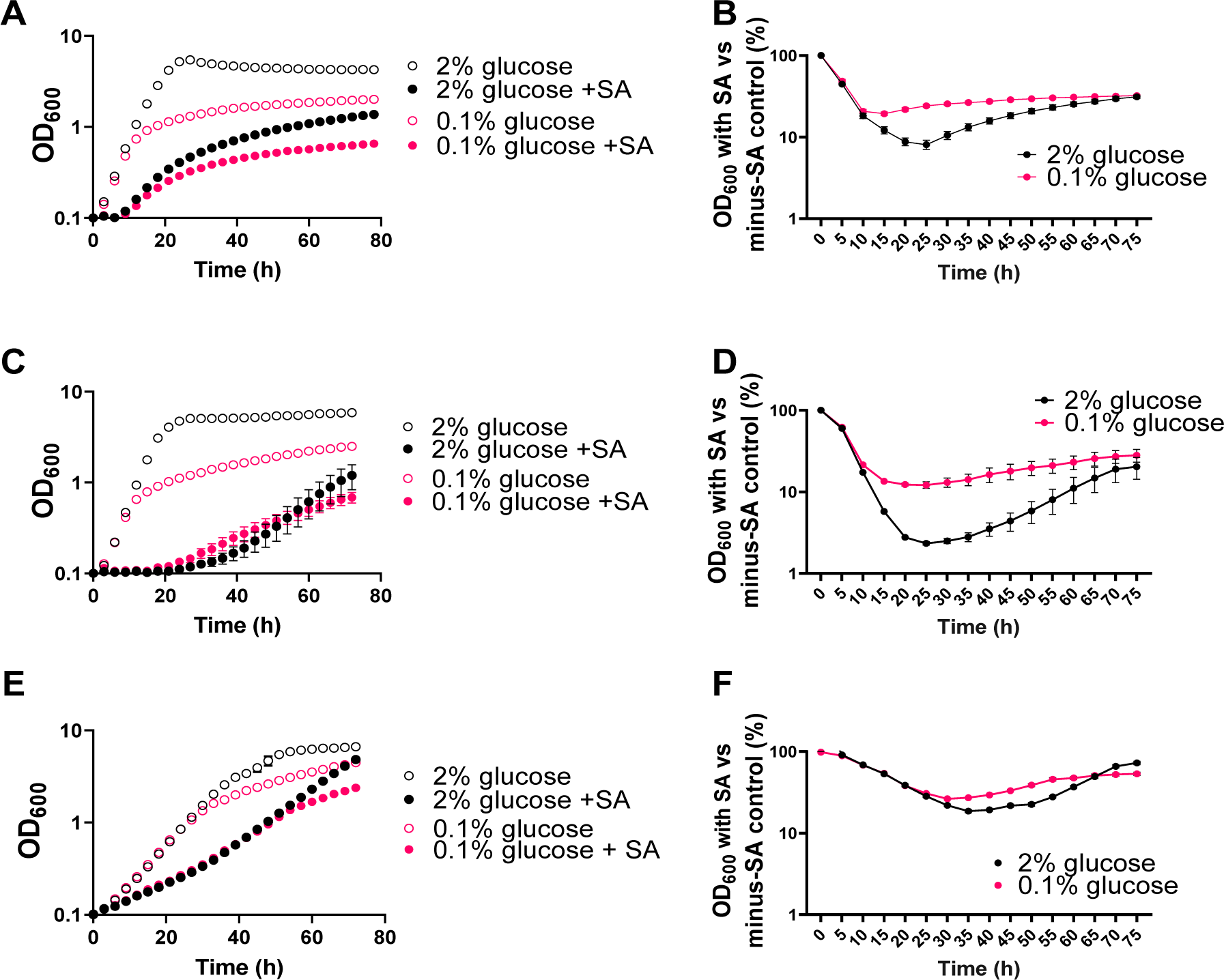
Spoilage yeasts show relative resistance to sorbic acid (SA) in low versus high glucose concentrations. Growth of *Z. bailii* (**A,B**), *S. cerevisiae* (**C,D**), and *B. bruxellensis* (**E,F**) is presented as raw OD_600_ values (A,C,E) or OD_600_ percentage versus no-sorbic acid controls (B,D,F) in 2% (black) or 0.1% (pink) glucose, either without (open symbols) or with (filled) sorbic acid. Sorbic acid was supplied at 2.5 mM, 1 mM and 2 mM for *Z. bailii, S. cerevisiae* and *B. bruxellensis,* respectively. Points represent means from three biological replicates; error bars (shown where they are larger than the dimensions of the symbols) represent SEM.

Other weak acids were tested for possible similar effects. In addition to sorbic acid, there was some relative resistance at low glucose to decanoic acid (Figure 2A,B), tested here because longer-chain acids have been shown to selectively inhibit respiring vs fermenting cells (Stratford et al., 2020). In contrast, a resistance phenotype at low glucose was not evident in cells cultured in the presence of acetic and propionic acids (Figure 2), shorter-chain acids that show little selective inhibition of cells growing by respiration (Stratford et al., 2020). The resistance phenotype was not dependent on the weak acid concentration as the effect was reproducible across a range of sorbic acid concentrations, even when very weakly inhibitory (Sup. Fig. 1).

**Figure 2.**
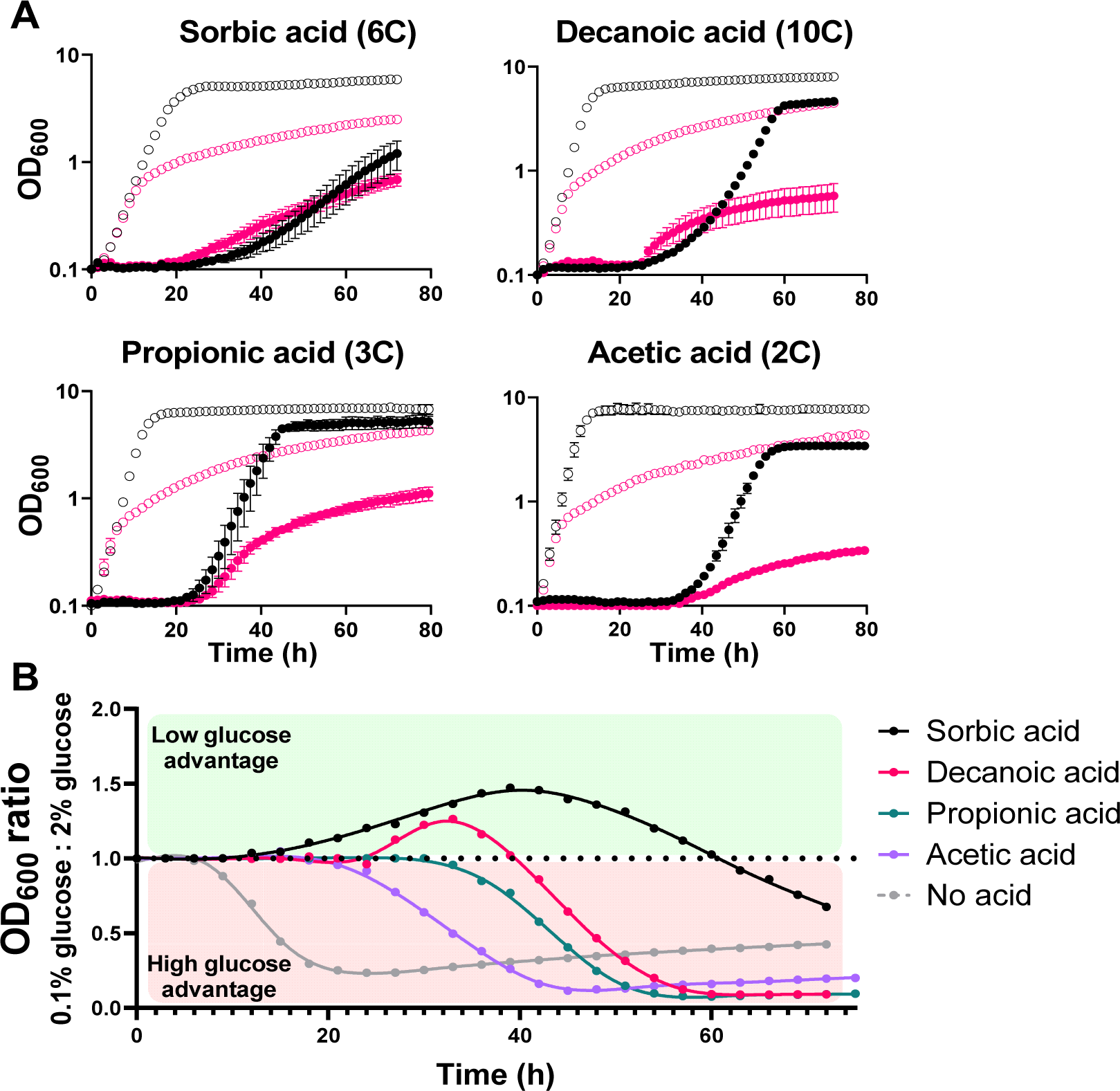
Growth in low glucose supports resistance to weak acids (WA) that selectively inhibit respiring cells. **A.** Growth of *S. cerevisiae* W303 in the presence of different WAs, with either 2% (black) or 0.1% (pink) glucose in the absence (open symbols) or presence (filled) of WA. Points represent means from three biological replicates. Carbon chain lengths are indicated after the relevant WA name. **B.** Growth data from A presented as the ratio of optical density in 0.1% versus 2% glucose cultures with each WA, where a ratio greater than 1.0 (above the dotted line) represents higher OD_600_ with the WA when in low glucose. Error bars (shown where larger than the dimensions of the symbols) represent SEM from three biological replicates.

Previous studies have implicated intracellular pH as a factor in WA resistance or sensitivity (Stratford et al., 2013, Stratford et al., 2014, van Beilen et al., 2014), while culturing with low glucose has been reported to acidify the cytosol of yeast cells (Elsutohy et al., 2017). To examine whether differences in internal pH could be related to WA resistance at low glucose, the intracellular pH of exponential-phase *S. cerevisiae* cells was measured with the pH dependent stain CFDA-SE (Stratford et al., 2014) and analysis by ratiometric flow cytometry, after 4 h of growth in either 2% or 0.1% glucose with acetic, sorbic, or decanoic acids (Sup. Fig. 2). Whereas intracellular pH was higher in low versus high glucose for cells incubated with decanoic acid, there was no significant effect of the glucose concentration with acetic acid, sorbic acid or without WA (Sup. Fig. 2). This suggested that the resistance phenotype at low glucose was not related to differences in intracellular pH.

Since sorbic acid has been shown to inhibit respiration, we also widened out the panel of yeast isolates to encompass yeasts reported to be fermentation deficient (Stratford et al., 2020), in order to determine the prevalence of the resistance phenotype when respiration was the predominant energy generating mechanism. Notably, we found that fermentation deficient yeasts typically did not show enhanced resistance at low glucose, with the phenotype present in only one (one of two *Rhodoturula glutinis* isolates) out of five fermentation deficient isolates tested (Sup. Fig 3). This observation suggested that fermentative metabolism may typically be required to elicit sorbic acid resistance in low glucose conditions.

To examine further a role for fermentation in sorbic acid resistance at low glucose, sorbic acid challenge experiments were conducted in anaerobic conditions, to force fermentative growth. Measurements of OD_600_ were taken at timepoints where the greatest difference in resistance between high and low glucose had been observed under aerobic conditions. In the anaerobic environment (inhibiting respiration but not fermentation), sorbic acid resistance in *S. cerevisiae* was enhanced in both of the glucose conditions compared to the aerobic environment (Figure 3). Furthermore, the difference between aerobic and anaerobic growth with sorbic acid was more than 50% greater at low glucose compared to high glucose. This indicated that sorbic acid resistance conferred by fermentative growth was greater for cells in low glucose.

**Figure 3.**
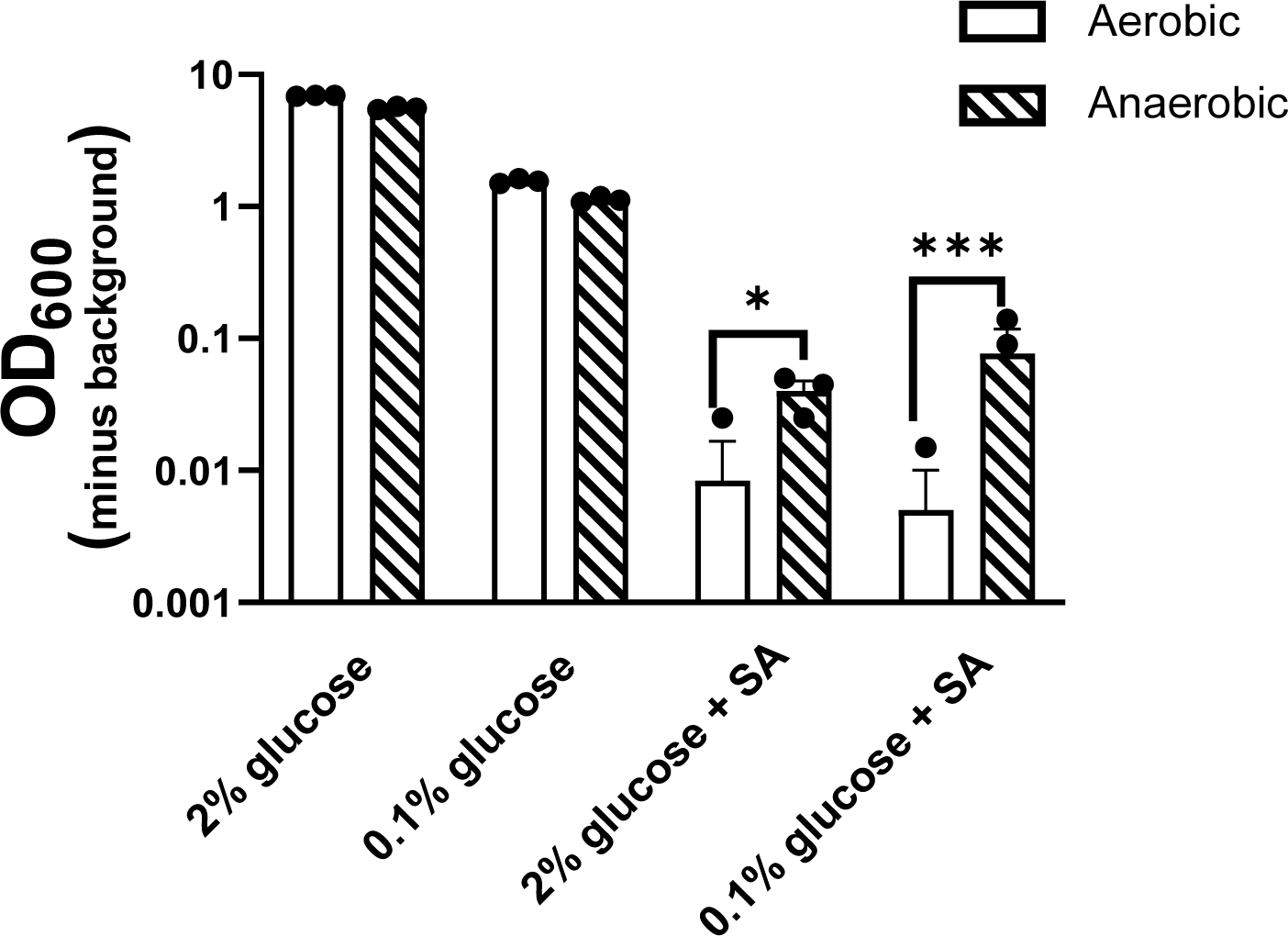
Growth in anaerobic conditions increases sorbic acid resistance, particularly at low glucose. Growth yields (OD_600_) of *S. cerevisiae* W303 were compared after 40 hours, static incubation at 24 °C in the different conditions with or without 1 mM sorbic acid (SA). Bars are means from three biological replicates (points). Where not all points are visible, absent points are OD_600_ value of zero and cannot be displayed on a logarithmic axis. Error bars represent SEM. * = p <0.05, *** = p < 0.001, according to one-way ANOVA.

To further probe the above findings, fermentation rate was measured according to pressure generated by gas evolution (Stratford et al., 2020) of cells in 2% or 0.1% glucose in the presence or absence of sorbic acid. Because yeast cells produce excess CO_2_ as a product of fermentation, whereas the net change in CO_2_ during respiration is balanced by O_2_ consumption, any increased gas pressure during yeast culture is predominantly the result of fermentation (apart from negligible amounts of other volatile molecules). Endpoint pressures determined after 24 h of growth in sealed bottles revealed that cells produced more gas (i.e., fermented more) with than without sorbic acid in both low and high glucose (Figure 4). This suggested that the presence of sorbic acid stimulates fermentative metabolism. A negative control of cells cultured with glycerol as carbon source, an obligatory respiratory substrate, showed no detectable fermentative activity. Notably, cells in low glucose showed little fermentation in the absence of sorbic acid, but more than a doubling in fermentative activity with the inclusion of sorbic acid. This compared with a <45% increase for the equivalent comparison in 2% glucose (Figure 4). This greater relative shift to fermentation at low glucose mirrored the greater relative resistance to added sorbic acid seen at low glucose (Figures 1–3) and the known resistance of fermentative cells to sorbic acid (Stratford et al., 2020). Consistent with this, growth on galactose (which can be fermented or respired, without the repression of respiration that occurs with glucose) did not give increased fermentation in the presence of sorbic acid and, likewise, did not give relative sorbic acid resistance at 0.1% versus 2% carbon source (Sup. Fig. 4).

**Figure 4.**
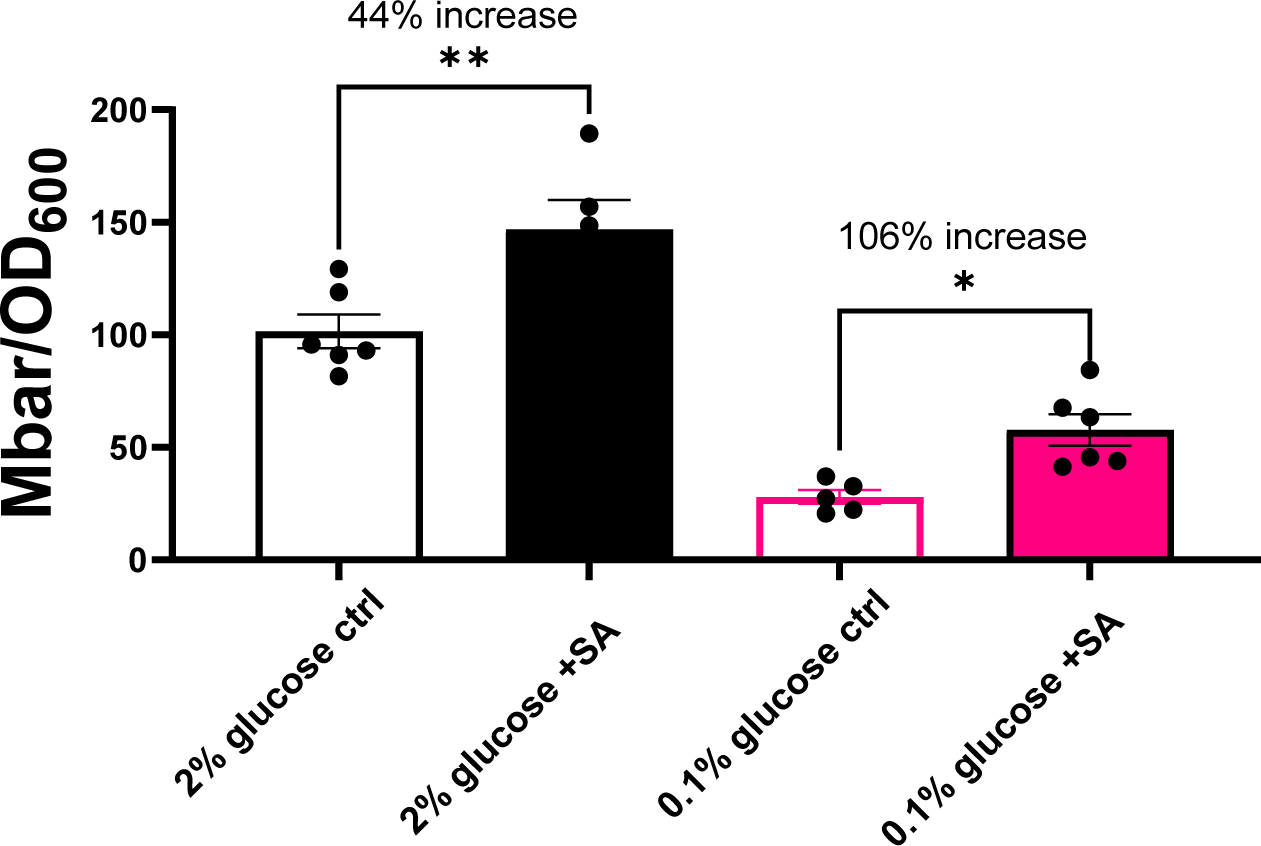
Sorbic acid increases fermentation by cells, particularly at low glucose. Pressure generated by *S. cerevisiae* W303 grown from starting inocula of OD_600_ 0.2 in the absence or presence of 1 mM sorbic acid in either 2% (black) or 0.1% (pink) glucose after 24 h static incubation at 24 °C, normalised by final OD_600_ readings. Error bars represent SEM from at least five biological replicates (points).

To pursue the hypothesis that a shift to increased fermentation is associated with sorbic acid resistance at 0.1% glucose, we sought to narrow down carbon sensing and metabolism pathways that may be integral to this phenotype. *S. cerevisiae* deletion strains lacking genes of the glucose sensing and repression pathways were assayed for retention of the relative sorbic acid resistance at 0.1% glucose seen in the wildtype. In total, 11 deletion strains were tested, as summarised in Figure 5 (see Sup. Fig. 5 for underlying data). Deletion of most of these genes had deleterious effects on sorbic acid resistance, both at high and low glucose. This was especially apparent with the deletion of *RGT2* (encoding a sensor of extracellular glucose), *YCK1* (kinase regulator of Rgt2), *MIG1* (transcriptional repressor of respiratory genes) and *REG1* (regulator of *MIG1*) (Figure 5C,D). Specific to the present report of low glucose induced sorbic acid resistance, deletion of *YCK1, MIG1,* or *RGT1* was more impactful in reducing sorbic acid resistance in low glucose conditions than in high glucose, implying that the relevant gene functions are indeed involved in sorbic acid resistance at low glucose.

**Figure 5.**
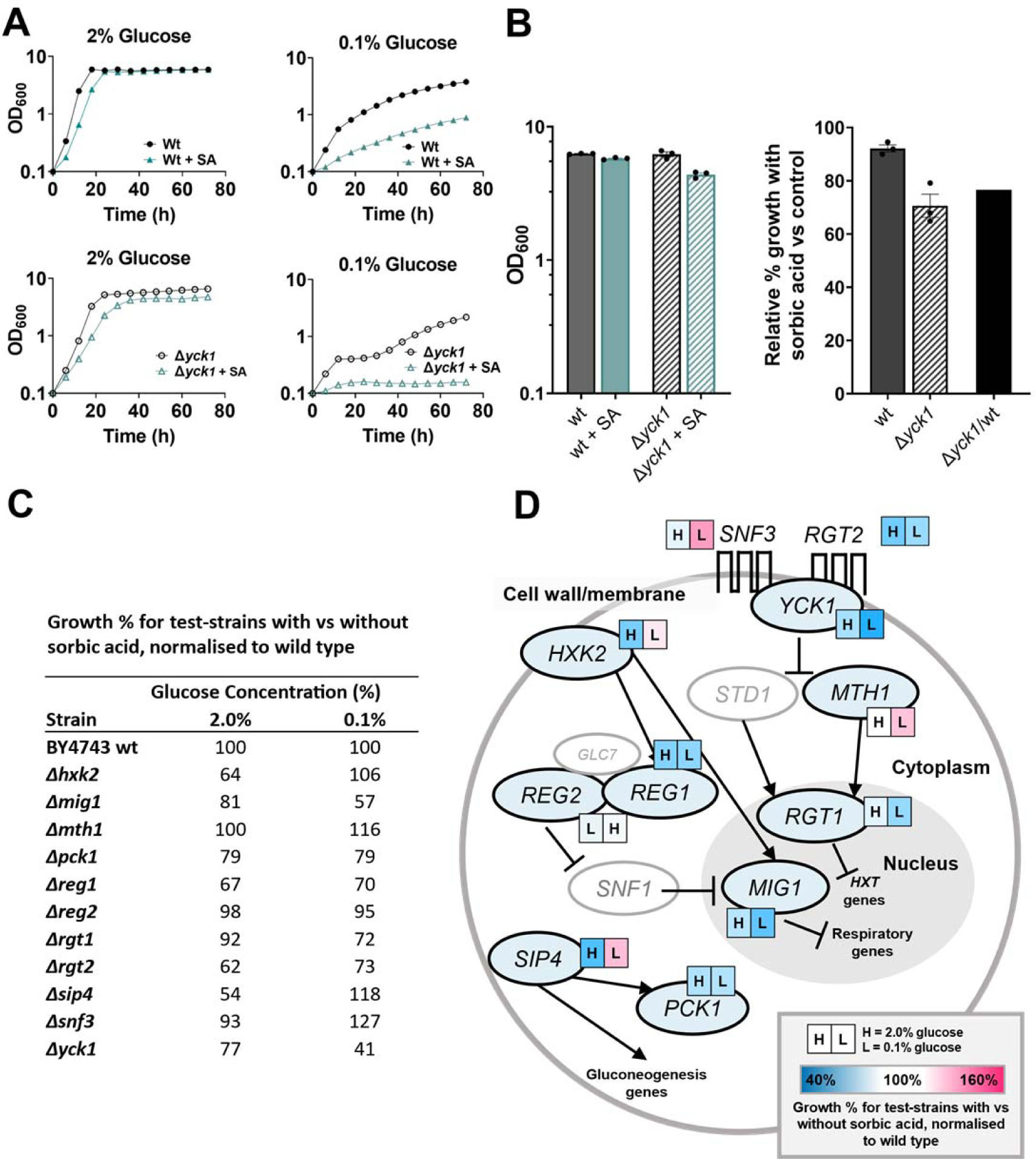
Deletion of key glucose sensing and repression genes alters sorbic acid (SA) sensitivity. **A.** Example growth assays for *S. cerevisiae* BY4743 (wt) and an isogenic deletion strain (example shown is Δ*yck1*), demonstrating sorbic acid hypersensitivity of the mutant at 0.1% glucose. Results from similar assays for other test-deletants are presented in Sup. Fig. 5. **B.** Example of data processing from growth assays in (A) to % inhibition values show in (C). Left panel: Wild type (solid) and *Δyck1* (hashed) OD_600_ at 60 h from the growth assays in 2% glucose either without (black) or with (green) 0.75 mM sorbic acid. Right panel: Using values from the left panel, growth of each strain with sorbic acid as a percentage of control growth (without sorbic acid). The final bar normalises the relative growth of the mutant with sorbic acid to that of the wild type and is the value used in panel (C). **C.** Growth of all tested deletants with versus without sorbic acid, expressed relative to the corresponding effect measured for the wild type, where values less than or greater than 100% represent relative sensitivity or resistance, respectively of the deletant to sorbic acid. **D**. Simplified scheme of the glucose sensing and signalling cascades examined in *S. cerevisiae* and phenotypes of corresponding deletion mutants. The results of (C) are presented here next to each gene as a colour scale for 2% (H) and 0.1% (L) glucose from blue to pink, with blue representing a sensitisation to sorbic acid and pink representing resistance, of the corresponding deletant relative to the wild type. Greyed-out components represent untested mutants.

In exception to the trend of decreased resistance in deletion strains, deletion of *SNF3* (a high affinity glucose sensor) or of downstream *MTH1* enhanced sorbic acid resistance in low glucose. Interestingly, Snf3 is a key sensor of low extracellular glucose concentrations, suggesting that (Snf3-dependent) sensing of low glucose environments mediates sorbic acid sensitivity, rather than the resistance we observed. Deletion of the transcription factor encoded by *SIP4*, a positive regulator of gluconeogenesis, also yielded sorbic acid hyper- resistance at low glucose but hyper-sensitivity in high glucose. However, deletion of *PCK1* encoding phosphoenolpyruvate carboxykinase, a key enzyme in gluconeogenesis, yielded only mild, negative impacts on sorbic acid resistance at both glucose concentrations.

The collective results, coupled with the evidence above that galactose (which does not directly interact with these signalling pathways) at low concentrations minimally induces sorbic acid resistance (Sup. Fig. 4), indicate that elements of upstream glucose sensing and glucose repression can be required for the relative sorbic acid resistance of cells at low glucose.

As increased fermentation appeared to be necessary for enabling sorbic acid resistance at low glucose, we hypothesised that encouraging respiratory metabolism could restore inhibition by sorbic acid. To achieve this, we supplemented cultures with succinic or malic acids, intermediate compounds of the tricarboxylic acid (TCA) cycle, during culture with sorbic acid. Supplementation of low glucose medium with succinic acid (a substrate of mitochondrial complex II; succinate dehydrogenase) both stimulated cellular respiration (Figure 6A), as also reported by Il’chenko et al. (2005), and markedly sensitized cells to sorbic acid (Figure 6B). Increased respiration in response to succinic acid supplementation was absent at 2% glucose (Figure 6A). Similar relative effects on growth were apparent also with malic acid supplementation at 0.1% glucose (Figure 6B), but these were weaker than with succinic acid; possibly because not all malic acid is used for succinate production via the TCA cycle and can instead feed into the malate-aspartate NADH shuttle (Easlon et al., 2008).

**Figure 6.**
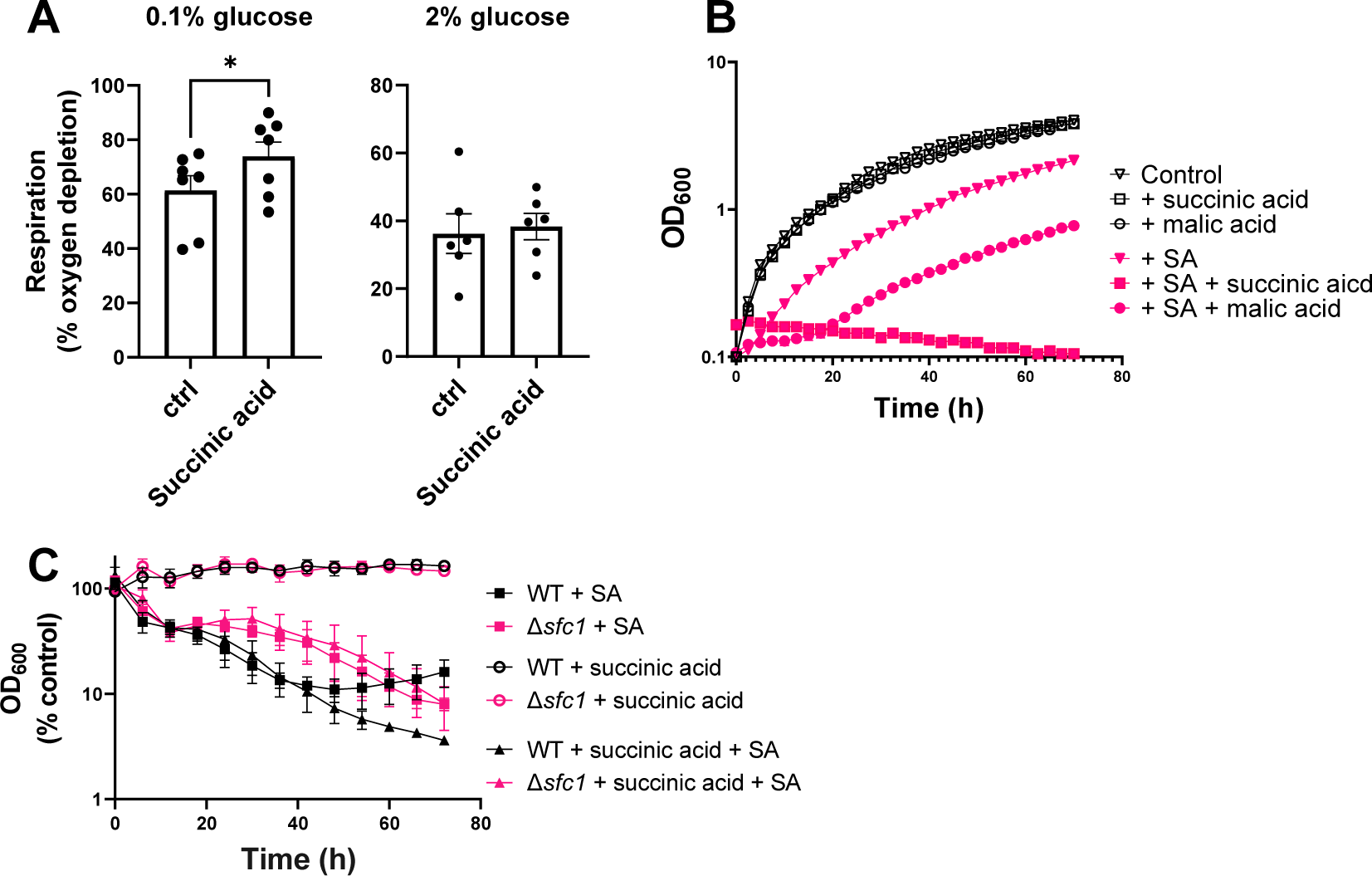
Supplementing yeast with succinic acid promotes respiration and sorbic acid hyper-sensitivity at low glucose. **A.** Measurements of respiration by *S. cerevisiae* W303 grown in the presence or absence of 111 mM succinic acid in YP supplemented with 0.1% glucose are presented as percentage oxygen depleted from culture medium after 1 h incubation at OD_600_ 0.5 (for time course of oxygen depletion, see Sup. Fig. 6). **B.** Growth curves with or without 1 mM sorbic acid and 111 mM of either succinic or malic acid are presented in 0.1% glucose. Growth curves are the average of three biological replicates; error bars (shown where they are larger than the dimensions of the symbols) represent SEM. (C) Growth of *S. cerevisiae* BY4743 (wt) and the isogenic *Δscf1* deletant in the presence of 1 mM sorbic acid, 111 mM succinic acid, or both, with OD_600_ presented as a percentage of values obtained at the corresponding timepoints for growth in medium without sorbic acid or succinic acid. Error bars represent SEM from three biological replicates.

In *S. cerevisiae*, succinate is transported into the mitochondria via the succinate-fumarate carrier (Sfc1) antiporter, and deletion of the *SFC1* gene prevents growth on respiratory substrates (Palmieri et al., 1997). To corroborate that succinic acid confers sorbic acid sensitivity via increased respiration, we conducted growth assays with the *S. cerevisiae Δsfc1* deletant. Results are presented as % growth versus the control condition without sorbic or succinic acid for easier visual comparison of the wild type and mutant. Sensitization to sorbic acid in the presence of succinic acid at 0.1% glucose became apparent only after around 40 h in the BY4743 wildtype background that is isogenic with the *Δsfc1* deletant (Figure 6C). Nevertheless, this succinic acid-dependent hypersensitivity was alleviated in the mutant lacking Sfc1 function.

Taken together, these findings offer not only a potential means of sensitizing spoilage yeasts to sorbic acid at low glucose, but they also support the hypothesis that a metabolic shift from respiration to fermentation enables yeast adaptation and resistance to sorbic acid resistance at low glucose.

## Discussion

Many food producers have been transitioning to reduced-sugar foods and beverages, in response to government legislation (in some countries) or consumer preferences (WHO, 2022). However, impacts of these changes on microbial preservative resistance and spoilage are barely characterised. The present results indicate that spoilage yeasts in low glucose are more resilient to inhibition by added sorbic-acid preservative than in high glucose. This finding contrasted with our starting hypothesis, which had been based on prior knowledge that: (i) sorbic acid targets yeast respiratory metabolism, evidenced during growth on a carbon source such as glycerol that can only be respired (Stratford et al., 2020); (ii) glucose scarcity favours respiratory over fermentative metabolism in many yeasts (Kayikci and Nielsen, 2015). The present findings offer an explanation for this apparent contradiction, as they indicated that cells in low glucose switch to fermentation in response to sorbic acid addition so enabling resistance. Other tested parameters like intracellular pH changes, which can be effected both by altered extracellular glucose levels (Martinez-Munoz and Kane, 2008, Orij et al., 2009) and weak acid exposure (Cole and Keenan, 1987, Stratford et al., 2013), were not correlated with sorbic acid resistance in the present conditions. In contrast, relative resistance to sorbic acid at low glucose was suppressed in certain deletants lacking key genes of the glucose signalling and repression pathway, consistent with this pathway playing a role in the (increased fermentation and) resistance response.

We propose that an ability to shift to a more fermentative metabolism in response to sorbic acid at low glucose promotes resistance as it mitigates the usual need for (sorbic acid-susceptible) respiration (Stratford et al., 2020) to support growth in this condition. Generally, cells must expend energy to mount an adaptive stress response, which requires the generation of ATP. In response to industrially relevant stressors (e.g. ethanol, salt, temperature), it has been demonstrated in yeast that respiration can be insufficient to meet these energy demands, and yeast cells can indeed switch to respiro-fermentative metabolism to mitigate the ATP deficit at the cost of biomass accumulation (Lahtvee et al., 2016). However, unlike in Lahtvee et al., (2016), where increased fermentation was coincident with decreased biomass accumulation, in the present study increased fermentation at low glucose with sorbic acid was correlated with increased biomass relative to the effect of sorbic acid at high glucose. This could reflect the additional respiration-targeting effect of sorbic acid compared to certain other stressors (i.e. decreased respiration alone helps protect against sorbic acid) or some other sorbic-acid specific element of the response. One effect of decreased respiratory activity would be decreased production of reactive oxygen species (ROS) from oxidative phosphorylation in the mitochondria (Suski et al., 2018). This is relevant here as it should limit ROS production by sorbic acid itself, an effect thought to lead to the depletion of functional respiratory complexes (i.e. respiratory inhibition) through petite-cell formation and FeS cluster targeting (Stratford et al., 2020).

It remains unclear how sorbic acid specifically might trigger such a switch in carbon metabolism. In the bacterium *Bacillus subtilis*, sorbic acid has been shown to promote responses similar to those seen during nutrient limitation, including upregulation of TCA- cycle associated genes (Ter Beek et al., 2008). However, results from a transcriptomic and proteomic study of sorbic acid exposure in *S. cerevisiae* indicated relatively little change in carbon metabolic or fermentation-associated gene expression that might also impact sorbic acid resistance (de Nobel et al., 2001). For example, whilst de Nobel et al. (2001) observed that expression of the glycolytic gene *TDH1* was upregulated in response to sorbic acid, deletion of the gene did not elicit any measurable change in sorbic acid resistance. However, it is important to note that this previous study was at 2% glucose and the importance of these or other genes up/down-regulated by sorbic acid at, say, 0.1% glucose may have greater impacts for sorbic resistance in the lower glucose condition examined in this study

As any increased preservative resistance of spoilage yeasts in reduced sugar foods could be a significant concern for the industry and consumers, we considered ways to mitigate the effect that could be feasible in real-world foodstuffs. Succinic acid is a metabolic intermediate of respiratory metabolism and a food-safe additive (Tretter et al., 2016). Stimulation of yeast respiration in the presence of succinic acid (at low glucose) in the present study was consistent with previous findings (Il’chenko et al., 2005). Moreover, the fact that this introduced bias to respiration was coincident with loss of sorbic acid resistance at low glucose suggested that such a feasible approach for metabolic re-routing may be sufficient for restoring preservative sensitivity of spoilage yeasts in low sugar formulations. That may be especially so as use of succinic acid supplementation as an effective means to increase yeast respiratory metabolism is not confined to the yeasts studied here (Il’chenko et al., 2005), suggesting the possibility of broad-spectrum intervention to help preserve low-sugar foods and beverages. Of course, any additions to foods (e.g., succinic acid) would require careful consideration also of potential flavour, texture or regulatory consequences.

As producers and consumers transition towards reduced-sugar food and drink formulations, efforts should be made to understand how these alterations change the spoilage-propensity landscape. The results of this study indicate that low sugar environments can facilitate preservative resistance by altering microbial carbon metabolism. However, reduced-sugar formulations often substitute sugar with low-calorie sweeteners, such as aspartame, acesulfame K, sucralose, and stevia (Lacroix et al., 2012). Although these ‘artificial sweeteners’ have little to no calorific value to humans, many can be metabolised by microorganisms (Lacroix et al., 2012), potentially providing an alternative energy source for spoilage organisms. Furthermore, additional formulation changes such as reducing acidity to accommodate reduced sugar or replacing the glucose in energy drinks with caffeine or B vitamins, may also constitute changes to the propensity for spoilage that warrant investigation.

## Acknowledgements

This work was supported by the Biotechnology and Biological Sciences Research Council (Grant Number BB/T014784/1).

## Supplementary Data

**Supplementary Figure 1.**
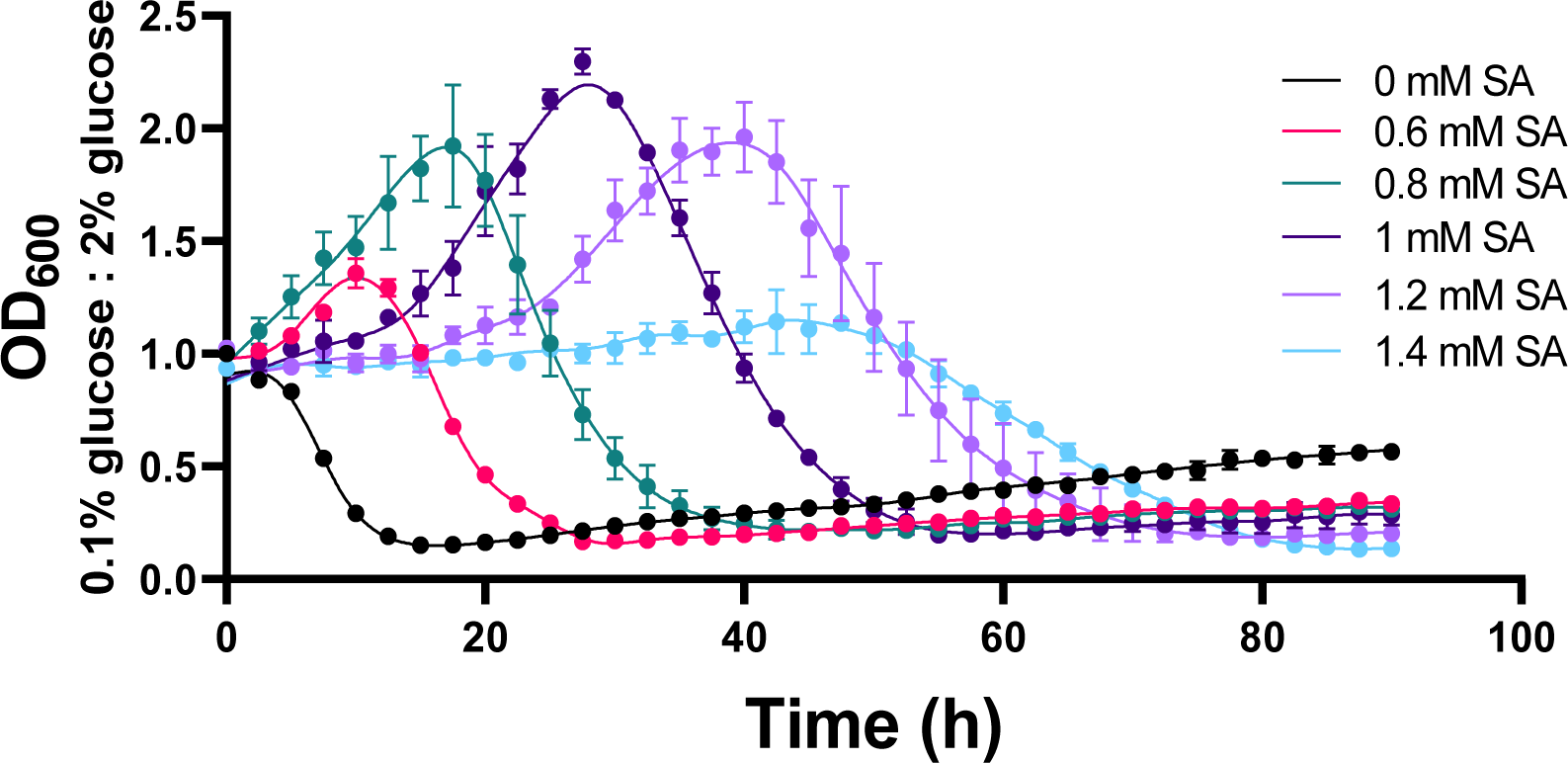
Relative sorbic acid resistance at low versus high glucose is evident over a range of sorbic acid concentrations. Growth data for *S. cerevisiae* with varying concentrations of sorbic acid (SA) are presented as the ratio of optical density in 0.1% versus 2% glucose cultures at each indicated time point of growth, where a ratio above 1.0 represents greater OD_600_ in the low glucose condition. Points represent the means from three biological replicates; error bars represent SEM of three biological replicates.

**Supplementary Figure 2.**
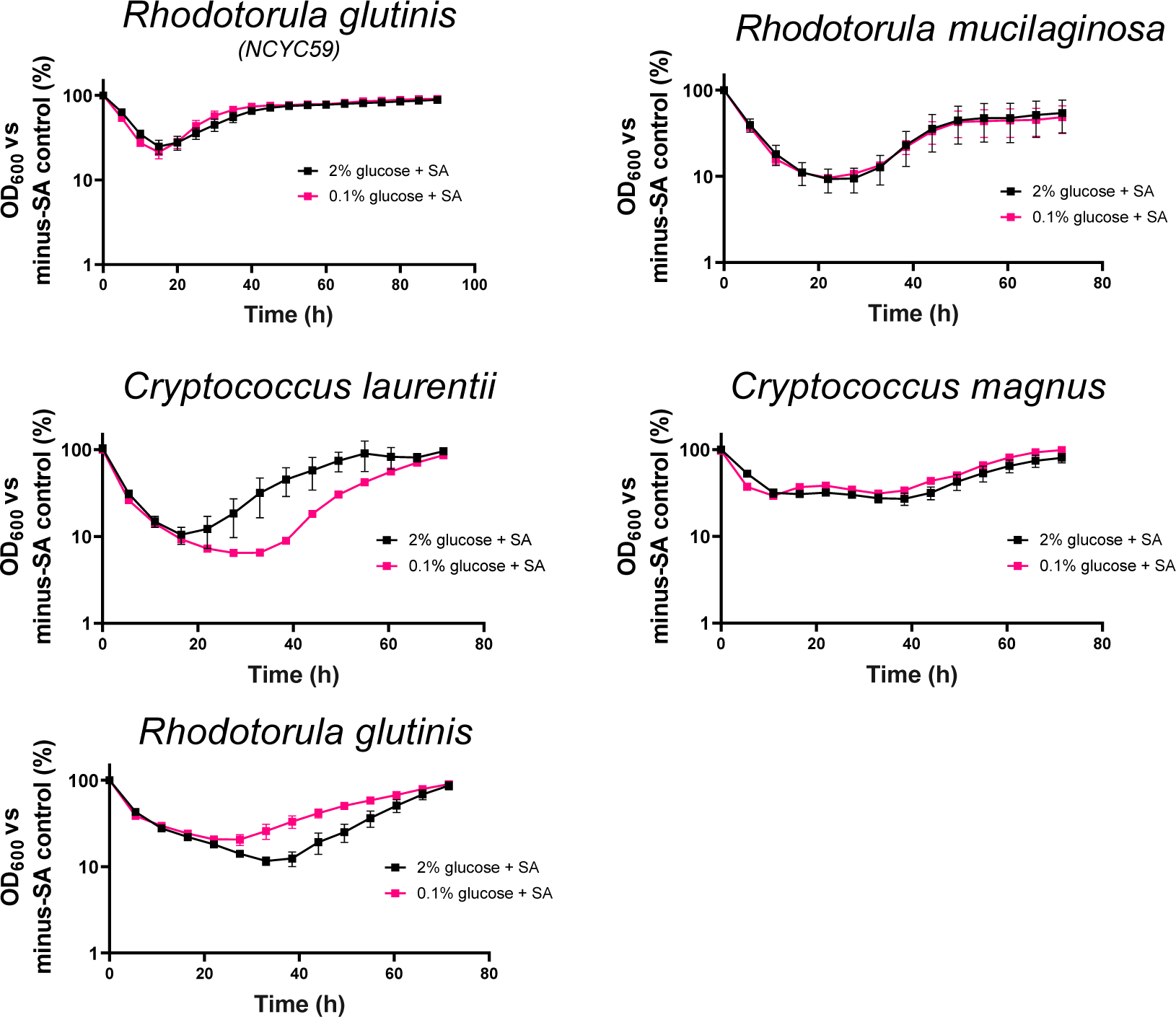
Intracellular pH of cells at different glucose levels, treated with weak acids. Intracellular pH of *S. cerevisiae* after 4 h incubation in YEP supplemented with either 2% or 0.1% glucose in the presence or absence of WAs (supplied at ∼55% of their MICs). Error bars represent SEM from three biological replicates, with each replicate value representing the median pH of 10^5^ cells.. ***, p < 0.001, according to one-way ANOVA; ns, not significant.

**Supplementary Figure 3.**
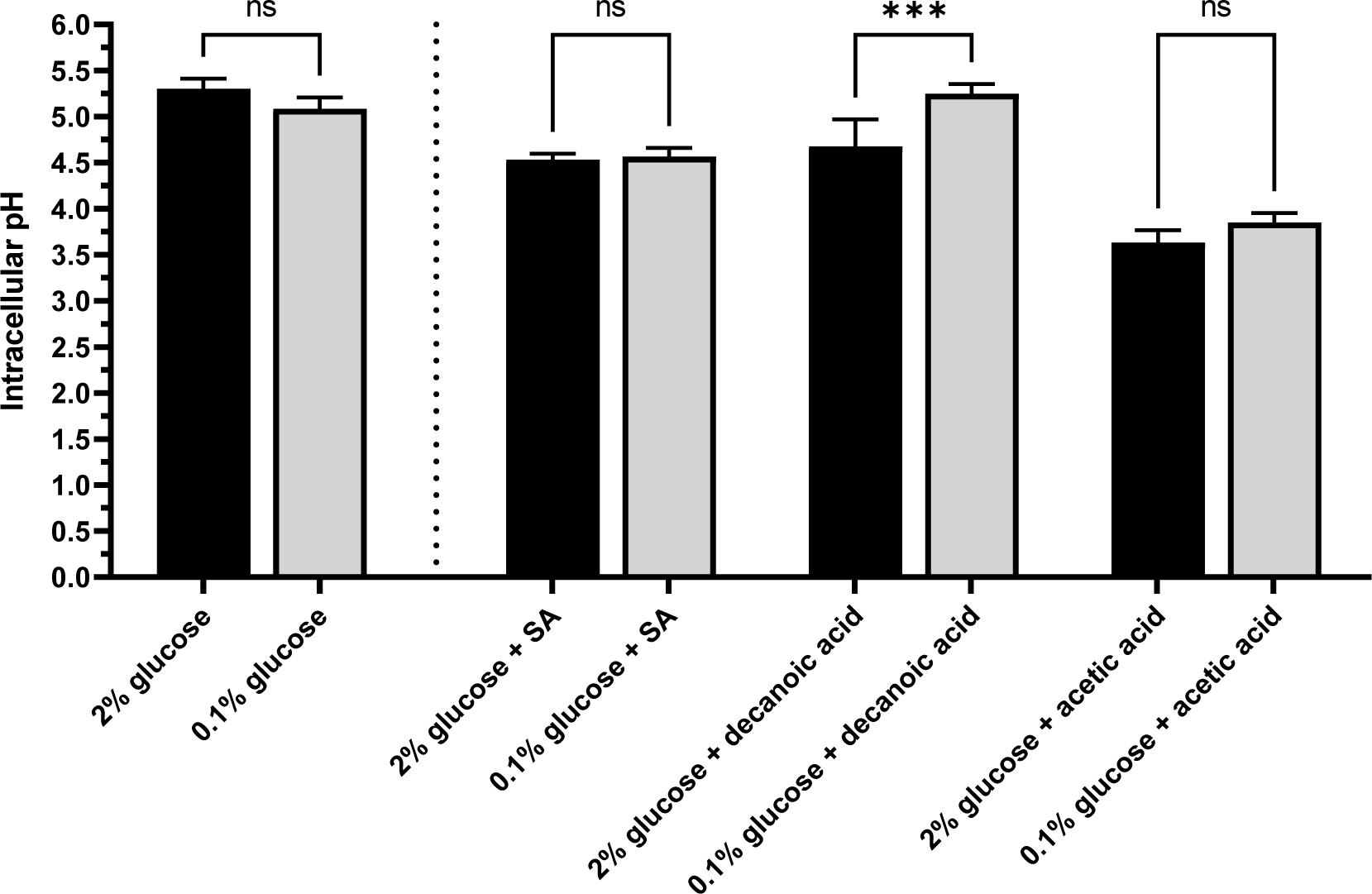
Absence of sorbic acid resistance at low glucose in several fermentation deficient yeasts. Growth curves for fermentation deficient yeast species in the presence of sorbic acid (SA; *R. glutinis* at 0.275 mM SA; *R. mucilaginosa,* 0.3 mM; P. *laurentii,* 0.4 mM; *C. magnus,* 0.1 mM) are represented by OD_600_ as a percentage of no-sorbic acid controls in 2% (black) or 0.1% (pink) glucose. Points represent means from three biological replicates, error bars (shown where larger than the dimensions of the symbols) represent SEM.

**Supplementary Figure 4.**
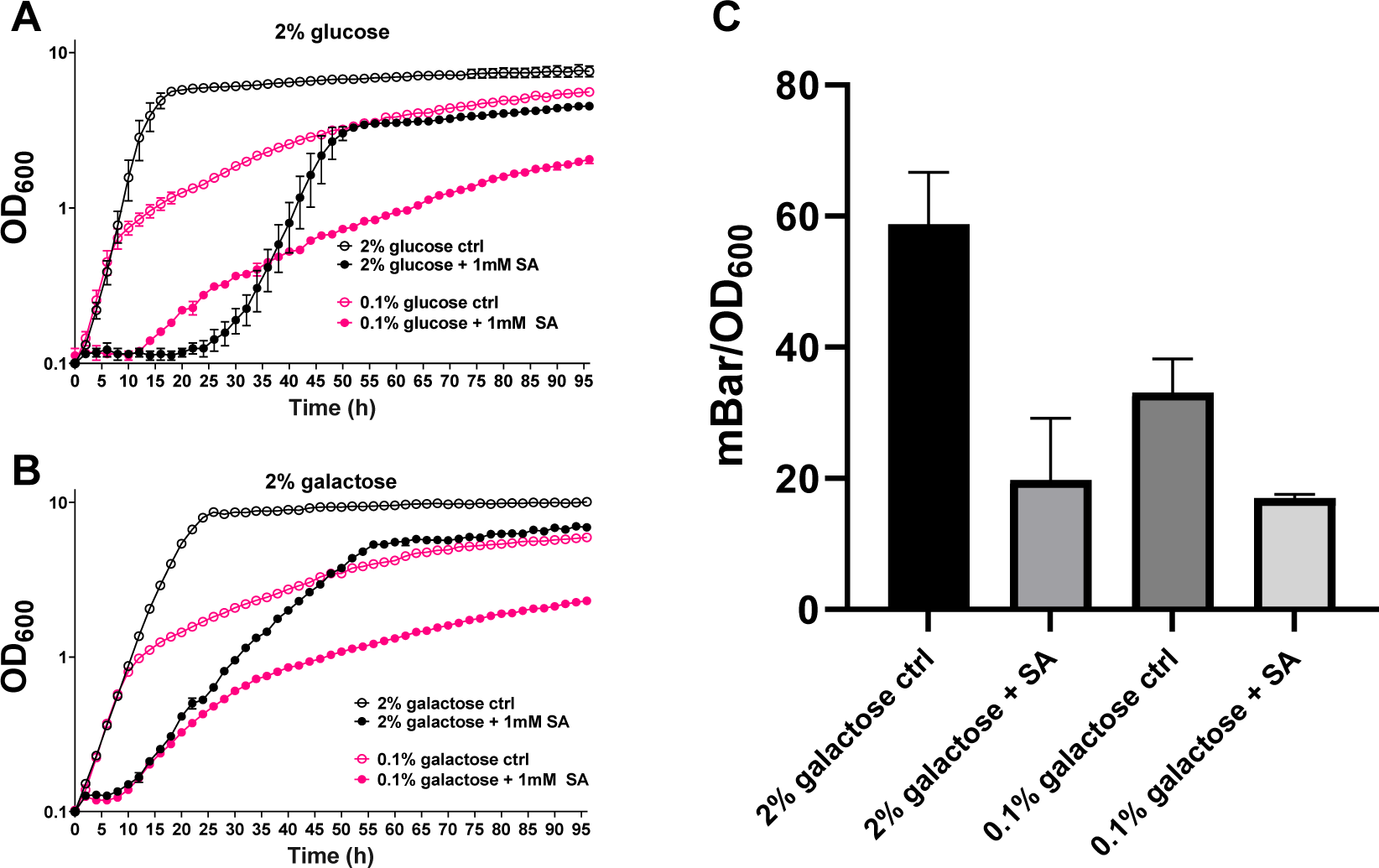
Effect of carbon source level on phenotype with sorbic acid (SA) depends on the carbon source. Growth curves for *S. cerevisiae* W303 grown with glucose (A) or galactose (B) at 2% (black) or 0.1% (pink) are presented as OD_600_ values either without (open circles) or with (filled circles) 1 mM sorbic acid. C. Pressure generated by *S. cerevisiae* W303 grown at a starting inoculum of OD_600_ 0.2 in the absence or presence of 1 mM sorbic acid in either 2% or 0.1% galactose after 24 h static incubation at 24 °C, normalised by final OD_600_ readings. Points represent the means of three biological replicates; error bars represent SEM.

**Supplementary Figure 5.**
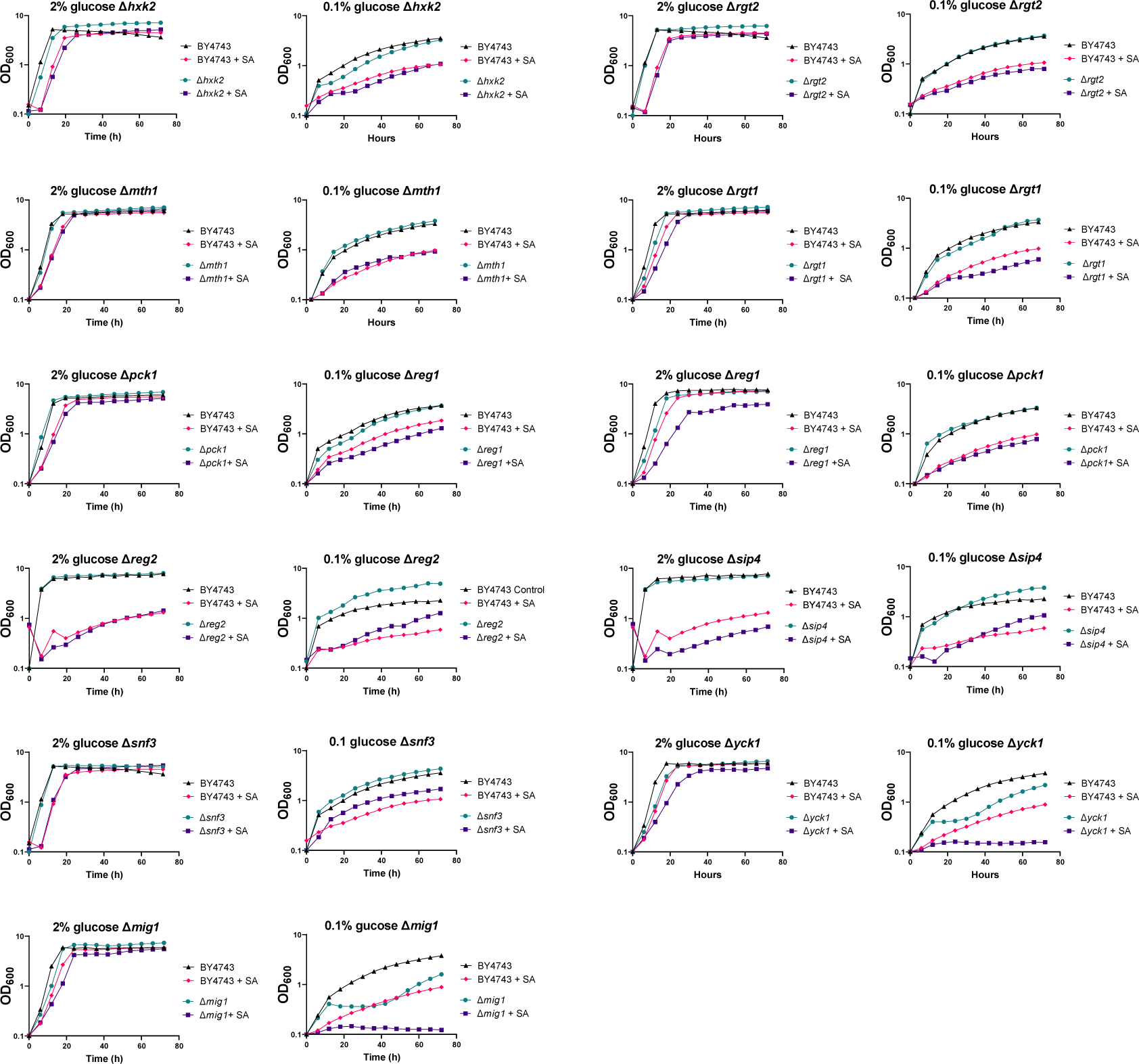
Growth assays with sorbic acid for deletion mutants relevant to glucose sensing and repression. Growth assays of *S. cerevisiae* BY4743 (wild type) and isogenic deletions strains at 2% and 0.1% glucose with or without 0.75 mM sorbic acid. Points represent means from three biological replicates, error bars (shown where larger than the dimensions of the symbols) represent SEM. Results are summarised in Figure 5 of the main manuscript.

**Supplementary Figure 6.**
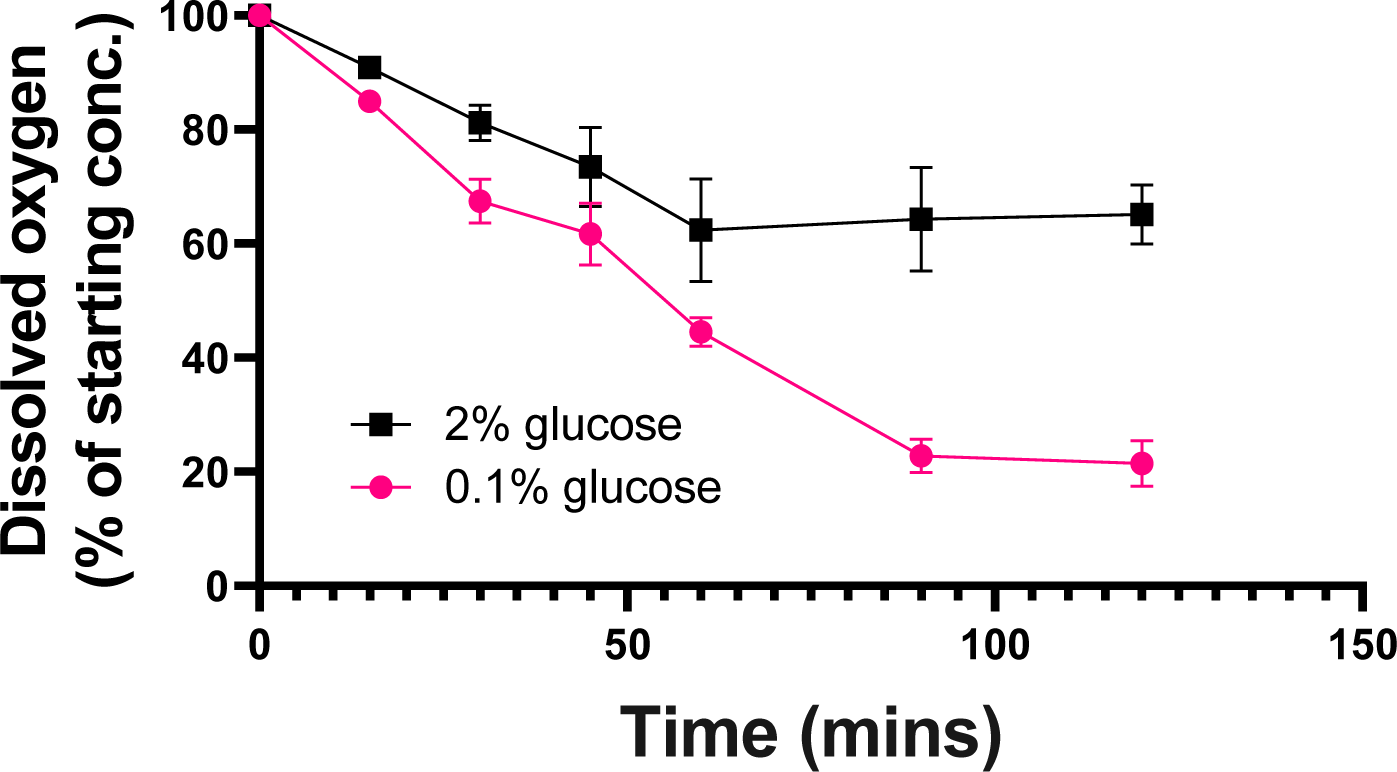
Time-course of oxygen depletion by *S. cerevisiae* W303. Cells were incubated in YP supplemented with either 2% (black square) or 0.1% (pink circle) glucose at an initial inoculum of OD_600_ 0.5. Oxygen concentration in medium is expressed as a % of starting concentration. Error bars represent SEM, N = 3.

